# Evaluation of the Prognostic Value of Four ECM-Associated Genes in Recurrence and Metastasis of Colorectal Cancer

**DOI:** 10.1101/2025.01.13.632700

**Authors:** Seçil Durel Avcıoğlu, Sulgun Charyeva, Beyza Doğanay Erdoğan, Özlem Kaymaz, Abdullah Durhan, Bülent Aksel, Ozan Yazıcı, Berna Savaş, Arzu Ensari, Hale Kıvrak, Evren Doruk Engin, Nevin Belder, Hilal Özdağ

**Affiliations:** Ankara University Biotechnology Institute; Ankara University Faculty of Medicine Department of Biostatistics; Ankara University Faculty of Science Department of Statistics; Ankara Training and Research Hospital; Etlik City Hospital; Gazi University Faculty of Medicine Department of Medical Oncology; Ankara University Faculty of Medicine Department of Medical Pathology; Etlik Zübeyde Hanım Gynecology Training and Research Hospital; Dokuz Eylül University Faculty of Medicine Department of Medical Microbiology

**Keywords:** Colorectal cancer, COL1A1, COL5A1, THBS2, FN1, Prognostic biomarker

## Abstract

Colorectal cancer is a significant public health issue due to its high incidence and high mortality rate, especially when diagnosed late. Identifying new biomarkers that can predict the diagnosis and prognosis of CRC is critical to reducing the mortality rate of the disease. This study tested the relationship of four extracellular matrix genes (COL1A1, COL5A1, THBS2 and FN1) with CRC prognosis at the mRNA and protein levels. Our previous studies identified these genes as candidate prognostic biomarkers for CRC. They were analysed in two cohorts: retrospective (130 CRC tumours and 70 healthy colon tissues) and prospective (160 paired tumours and normal tissues with 180 CRC serum samples).

Accordingly, in the retrospective cohort, COL1A1 (p>0.0001), COL5A1 (p>0.0001), THBS2 (0.0001) and FN1 (0.001) were expressed at higher levels in tumours compared to controls by qRT-PCR and in metastatic cases compared to non-metastatic cases. In the same cohort, immunohistochemistry analysis showed that increasing protein levels of COL1A1 and FN1 correlated with mRNA levels. In addition, high COL1A1 mRNA levels were associated with OS (p>0.011), RFS (p>0.0001), and DMFS (p>0.0001), high COL5A1 mRNA levels were associated with OS (0.042), RFS (0.013), DMFS (p>0.003) and high FN1 levels were associated with DMFS (p>0.039). In the prospective cohort, COL1A1, COL5A1 and THBS2 were found to be higher in tumour tissue compared to matched normal tissue and in metastatic tumours compared to primary tumours by qRT-PCR in 160 tumours and matched normal tissues. In this cohort, FN1 levels were unexpectedly higher in normal tissues than in tumour tissues. In the prospective cohort, where 180 CRC serum samples were also tested by ELISA, COL1A1 (p>0.0001), COL5A1 (p>0.05), and THBS2 (p>0.05) levels were higher in metastatic cases than in the sera of non-metastatic cases. As a result, this study associated COL1A1, COL5A1 and THBS2 genes, which have different data on their prognostic effects in the literature, with poor prognosis of CRC at both mRNA and protein levels.

## INTRODUCTION

The high prevalence and mortality rates of colorectal cancer (CRC), coupled with the limitations of existing diagnostic and prognostic tools, underscore the urgent need for novel prognostic biomarkers. CRC remains a leading cause of cancer-related deaths worldwide, accounting for nearly 900,000 deaths annually and ranking as the fourth most deadly malignancy [1]. A significant proportion of patients are diagnosed at advanced stages, severely limiting treatment options and survival rates [2–4]. This inadequacy highlights the need to develop novel biomarkers to improve patient stratification, guide treatment decisions, and ultimately enhance clinical outcomes.

The complex tumour microenvironment in CRC further complicates biomarker discovery. Inflammatory processes and angiogenesis play critical roles in tumour development and metastasis, and biomarkers related to these pathways are being actively explored for their prognostic value [5, 6].

Furthermore, the biological heterogeneity of CRC necessitates the identification of biomarkers that can accurately reflect individual tumour characteristics and predict patient responses to therapy. Integrating machine learning and bioinformatics approaches into biomarker discovery is a valuable strategy. This approach enables the identification of novel biomarkers by analysing large datasets [7, 8].

The extracellular matrix (ECM) plays a crucial role in the progression and metastasis of cancer [7]. Numerous studies have demonstrated that the composition and organisation of the ECM are significantly altered in colorectal cancer compared to healthy colon tissue [8–10]. These changes actively reshape the tumour microenvironment, driving key processes such as cell proliferation, migration, invasion, and angiogenesis. Beyond its functional role in tumour biology, the ECM emerges as a promising reservoir of potential biomarkers, offering new avenues for advancing CRC diagnosis and treatment [9].

These alterations in the ECM contribute to the development and progression of CRC by influencing various cellular processes, including cell proliferation, migration, invasion, and angiogenesis. Consequently, the ECM represents a rich source of potential biomarkers for CRC.

COL1A1, COL5A1, and FN1, as components of the ECM, and THBS2, a homotrimeric glycoprotein involved in cell-to-cell and cell-to-matrix interactions, are known to be dysregulated during carcinogenesis.

Elevated expression levels of COL1A1 have been consistently associated with poor prognosis in multiple cancer types, suggesting its utility as a prognostic marker. In colorectal cancer, COL1A1’s role as a prognostic biomarker is well-documented. Studies have shown that high COL1A1 expression is significantly associated with serosal invasion and lymphatic metastases [10, 11] and noted that COL1A1 expression levels could predict relapse in high-risk stage II and III colorectal cancer patients, emphasising its potential utility in clinical decision-making. Moreover, COL1A1 has been implicated in the epithelial-mesenchymal transition (EMT), a critical process in cancer metastasis, further solidifying its role in cancer progression [12].

COL5A1 is differentially expressed in several cancer tissues compared to normal tissues [13, 14]. Furthermore, COL5A1 expression levels have been associated with various clinicopathological parameters, including tumour size and lymph node involvement, reinforcing its relevance in disease progression. [13, 14]

In addition to its role in cancer progression, FN1 has been identified as a potential diagnostic marker. Studies have reported that FN1 levels in serum or tissue samples can indicate tumour presence and aggressiveness. For instance, circulating tumour cells expressing FN1 have been linked to poor prognosis in nasopharyngeal carcinoma, suggesting that FN1 could be utilised in monitoring disease progression [15]. Moreover, FN1 expression in tumour tissues has been correlated with various clinicopathological features.

Several studies have highlighted the role of THBS2 in CRC diagnosis and prognosis, particularly concerning metastatic potential and recurrence [16, 17]. Some studies indicate that high THBS2 expression correlates with poor prognosis and increased metastasis in CRC patients [16, 18]. Conversely, other research suggests that elevated THBS2 levels may not uniformly predict adverse outcomes across different patient cohorts or cancer stages [19, 20]. This inconsistency highlights a critical gap in understanding the precise role of THBS2 in CRC progression and its potential as a reliable prognostic marker.

Despite the promising implications of COL1A1, COL5A1, FN1, and THBS2 as biomarkers, the specific investigation of tumour protein expression and serum levels in colorectal cancer patients still requires further exploration. Current literature primarily focuses on tissue mRNA expression rather than immunohistochemical analysis and serum concentrations. Elucidating a possible correlation between tissue protein expression and circulating protein levels of these potential biomarkers may pave the way to assess the clinical utility of these proteins’ serum levels as non-invasive biomarkers for colorectal cancer.

Therefore, this study aimed to initially investigate the mRNA and protein expression of COL1A1, COL5A1, FN1, and THBS2 in tumour tissue in formalin-fixed paraffin-embedded (FFPE) sections of a retrospective cohort of CRC patients. Then, we analysed the mRNA expression level of these four genes in frozen section biopsies and paired normal tissue sections in a prospective CRC cohort along with their protein levels in corresponding serum samples.

## RESULTS

### COL1A1, COL5A1, FN1 and THBS2 mRNA Expression Levels in Colorectal Cancer (CRC) Tissue and their Association with Metastasis

In our previous study (Belder N et al., 2004, PeerJ under review), we showed the association of COL1A1, COL5A1, FN1 and THBS2 genes with the prognosis of colorectal cancer on 49 primary CRC tumours and matched normal tissues. We validated our results first with three independent GEO datasets, including 251 CRC tumours and 50 normal colon tissues and then with an independent cohort, including 64 CRC tumours and matched normal tissues. Validation of a biomarker candidates in independent patient cohorts is critical to ensure its robustness, reproducibility, and clinical applicability across different populations and conditions. Here, in this study, we demonstrated that these four genes were overexpressed in tumour tissues compared to normal tissues in 130 CRC tumours and 70 normal colon tissues, including the 64 normal tissues we analysed in our previous study (Table 1) (Fig 1 A, B, C, D Left side panels). On the other hand, it was also shown that these four genes were expressed at higher levels in patients who metastasised during the follow-up in the same discovery (retrospective) cohort (Fig 2. A, B, C, D Left side panels).

**Table 1.**
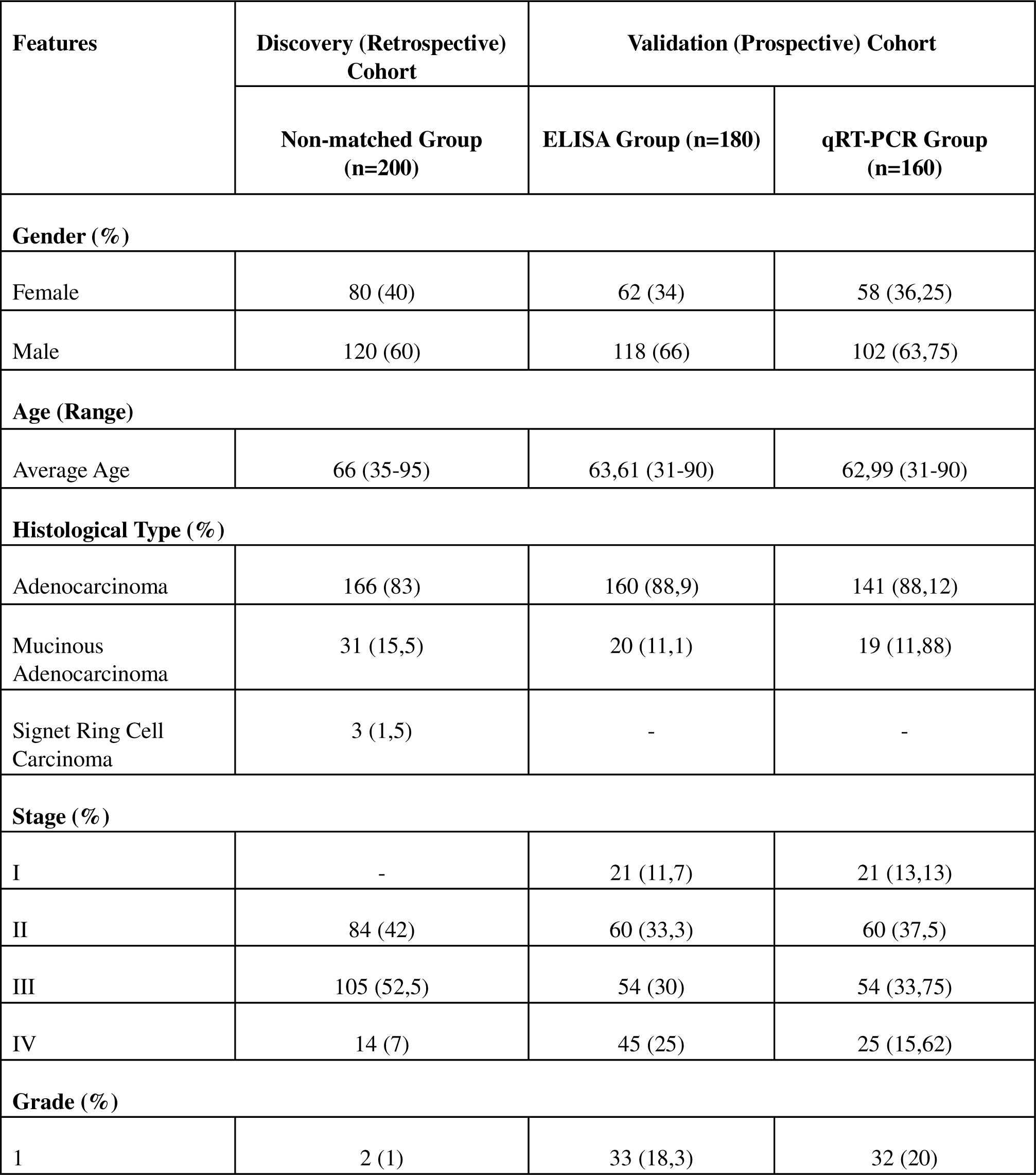
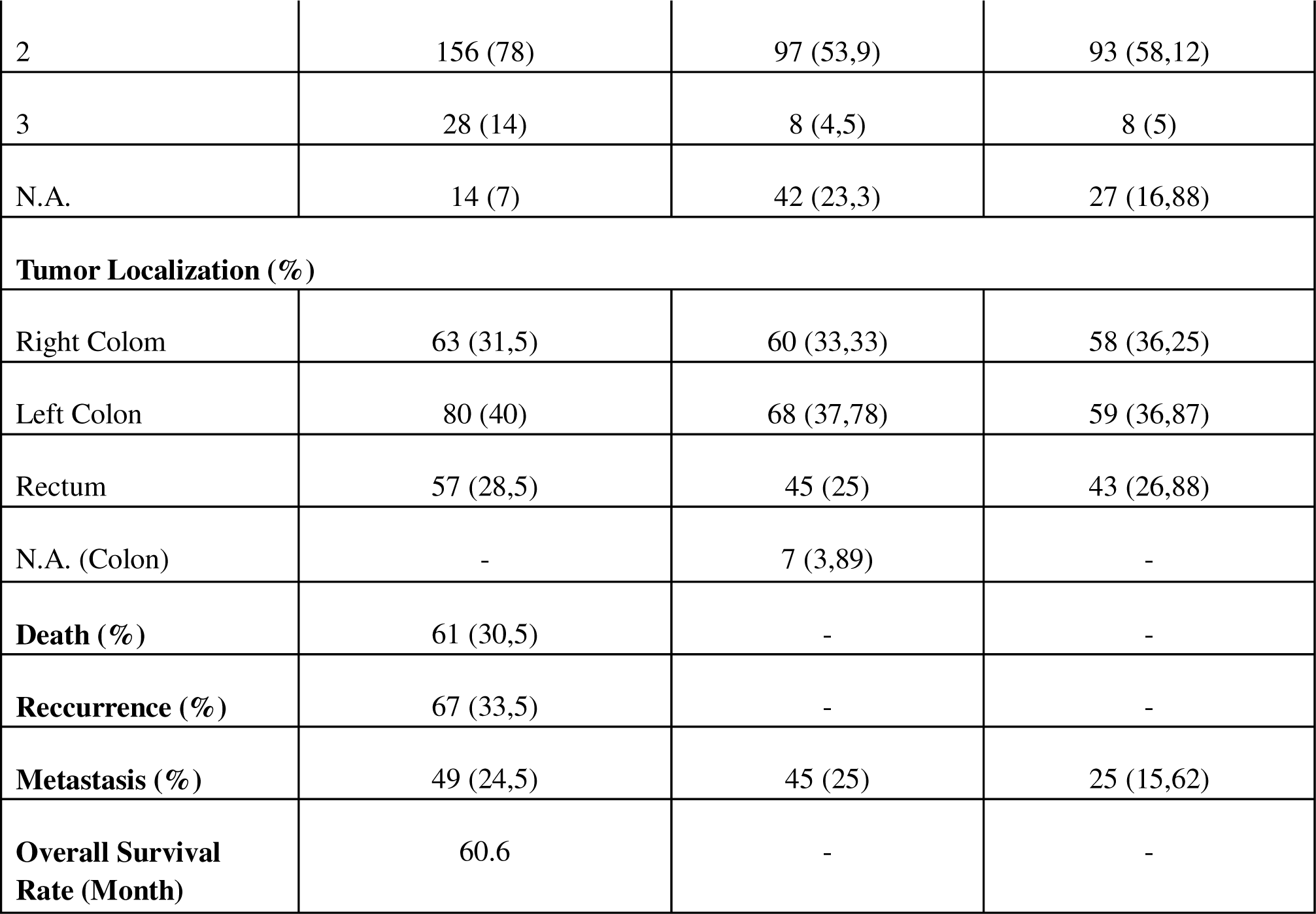
Clinicopathological characteristics of patients in the discovery (retrospective) and validation (prospective) cohorts.

**Figure 1.**
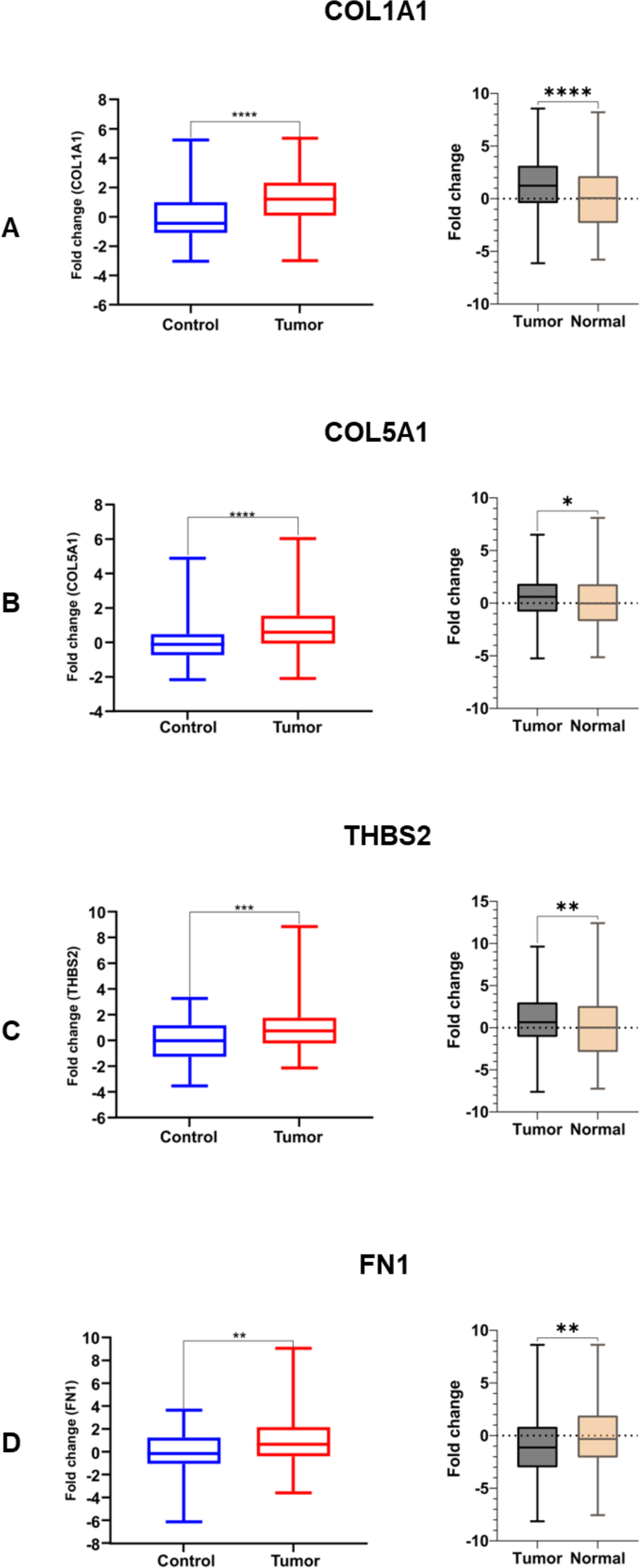
Colorectal tissue mRNA expression levels in the retrospective (discovery) and prospective (validation) cohorts. Left panels: Retrospective cohort. Right panels: Prospective cohort. A. COL1A1, B. COL5A1, C. THBS2, D. FN1.

**Figure 2.**
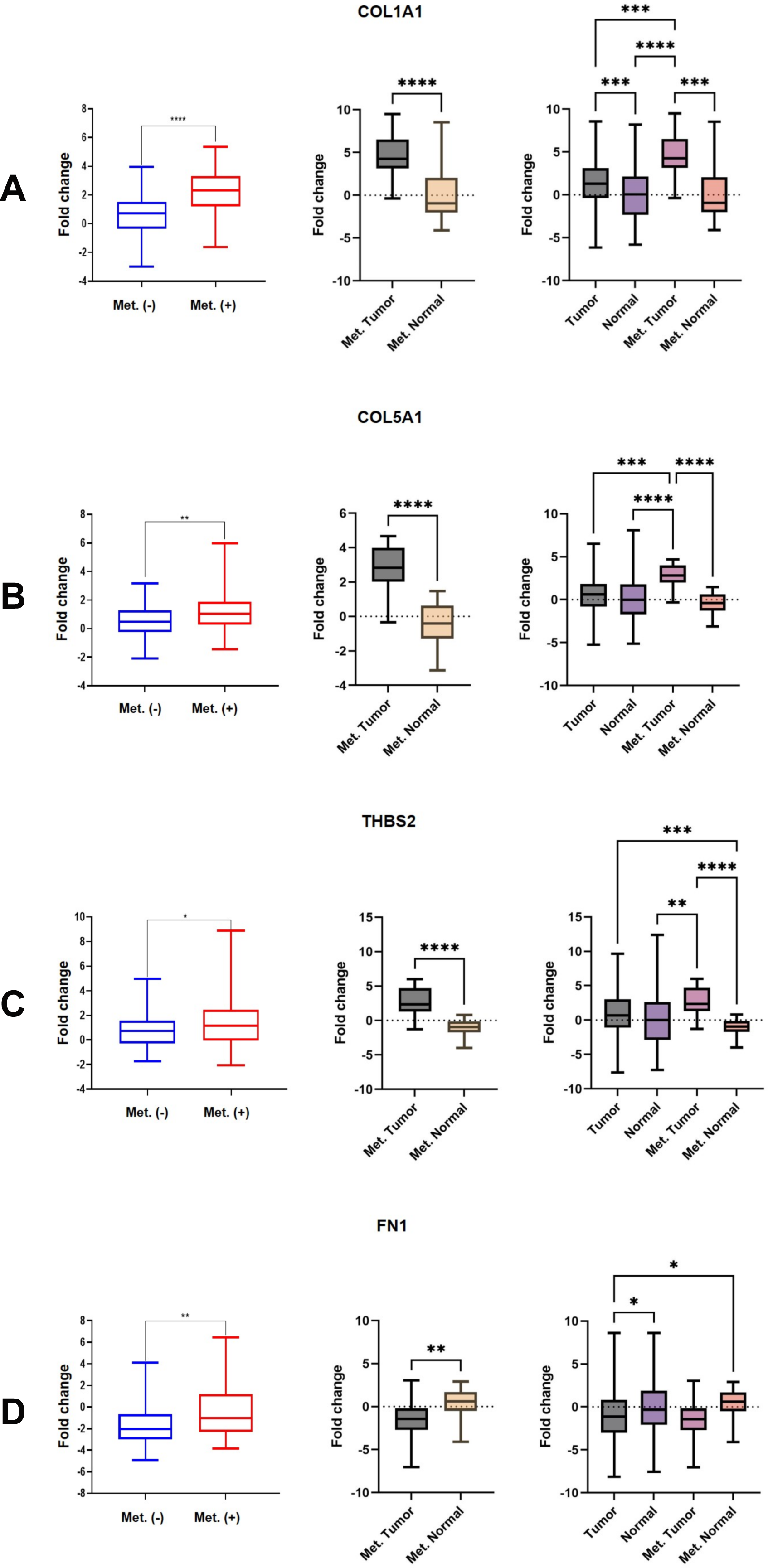
Colorectal tissue mRNA expression levels in the retrospective (discovery) and prospective (validation) cohorts between metastatic and non-metastatic cases. Left panels: Retrospective cohort. Middle panel: Primary tumour vs metastatic tumour. Right panel: Normal tissue vs. metastatic tumor. A. COL1A1, B. COL5A1, C. THBS2, D. FN1.

In the validation (prospective) cohort (Table 1), which was created to confirm the prognostic values of these four genes whose association with CRC prognosis was sought to be revealed, COL1A1, COL5A1 and THBS2 genes were shown to be expressed at higher levels in tumour tissues compared to matched normal tissues, based on the evaluation of 160 primary tumours (Fig 1, A, B, C right side panels). It should also be noted that the expression level of COL1A1, COL5A1 and THBS2 is statistically significantly higher in metastatic tumours than in primary tumours (Fig 2. A, B, C Right side panels). Unexpectedly, the FN1 gene was shown to be expressed at lower levels in tumour tissues compared to matched normal tissues and in metastatic tumours compared to matched normal tissues (Fig 1D and Fig 2D).

### COL1A1, COL5A1, FN1 and THBS2 mRNA Expression Levels in Colorectal Cancer (CRC) Tissue and their Association with Recurrence ve Survival

Since the retrospective discovery (retrospective) cohort in this study also included recurrence and survival information, the association between the tissue mRNA expression levels of the four genes in question with recurrence and survival was also examined. Accordingly, the expression of COL1A1 and COL5A1 genes was statistically significantly higher in patients who died than in those who survived (Fig 3 A, B first column). At the same time, there was no such relationship in the expression of THBS2 and FN1 genes (Fig 3 C, D first column). On the other hand, the expression of these four genes was higher in patients with recurrence than in those without (Fig 3 A, B, C, D second column).

**Figure 3.**
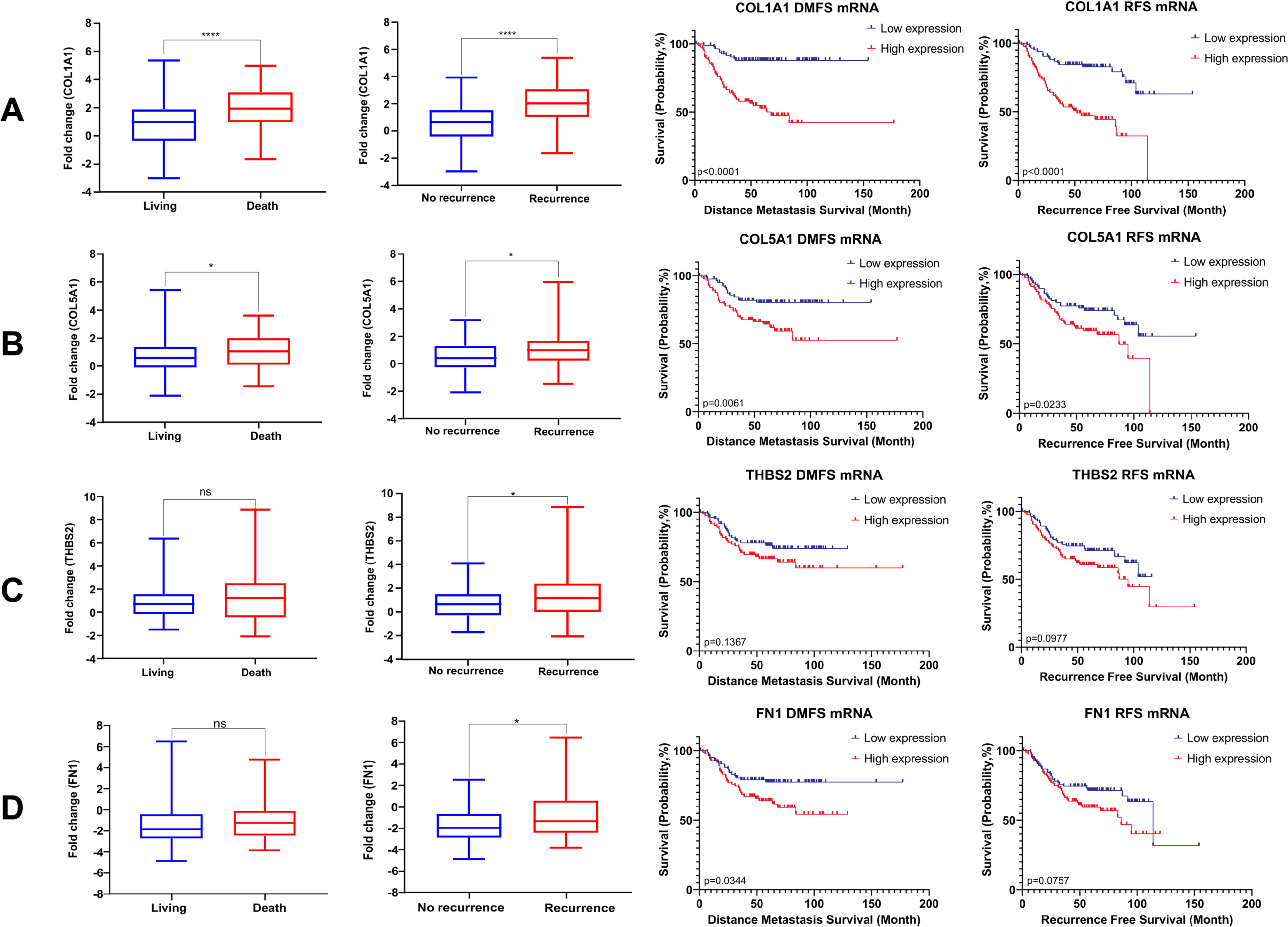
Association of COL1A1, COL5A1, THBS2 and FN1 mRNA expression levels with overall survival (OS), recurrence-free survival (RFS) and distant metastasis-free survival (DMFS) in the retrospective (discovery) cohort. A. COL1A1, B. COL5A1, C. THBS2, D. FN1.

Recurrence-free survival analyses in the discovery cohort indicated that high COL1A1 and COL5A1 gene expression reduced the survival rate in patients with recurrence (p<0.0001 for COL1A1 and p<0.013 for COL5A1) (Fig 3 A, B third column). At the same time, there was no association between THBS2 and FN1 gene expression and recurrence-free survival (Fig 3 C, D third column). On the other hand, high COL1A1, COL5A1 and FN1 gene expressions were shown to reduce the survival rate in distant metastasis-free survival analyses (p<0.0001 for COL1A1, p<0.003 for COL5A1, and p<0.039 for FN1) (Fig 3. A, B, D fourth column). There was no association between THBS2 gene expression and distant metastasis-free survival (p<0,13) (Fig. 3. C).

### COL1A1, COL5A1, FN1 and THBS2 Protein Expression Levels in Colorectal Cancer (CRC) Tissue and their Association with the Disease Prognosis

Immunohistochemical (IHC) staining was performed for COL1A1, COL5A1, and FN1 proteins on a cohort of 200 formalin-fixed, paraffin-embedded CRC tumour samples. To optimise staining efficiency and tissue usage, COL5A1 and FN1 were stained on single slides with two samples per slide, while COL1A1 was evaluated using tissue microarray blocks. Despite multiple attempts, we could not achieve a satisfactory staining pattern for THBS2 and therefore excluded it from further analysis.

COL1A1 IHC staining was assessed semiquantitatively by a pathologist, with 168 cases classified as negative (score 0) and 32 cases as positive (score 1). Positive staining was characterised by supranuclear punctate cytoplasmic staining. The presence of positive internal controls confirmed the efficacy of the staining procedure. As shown in Figure 4A (left and middle panels), a significant positive correlation was observed between COL1A1 protein expression (IHC score) and mRNA expression levels. Furthermore, survival analysis revealed that patients with positive COL1A1 staining exhibited significantly shorter overall, recurrence-free, and metastasis-free survival than those with negative staining (Figure 4A, right panel).

**Figure 4.**
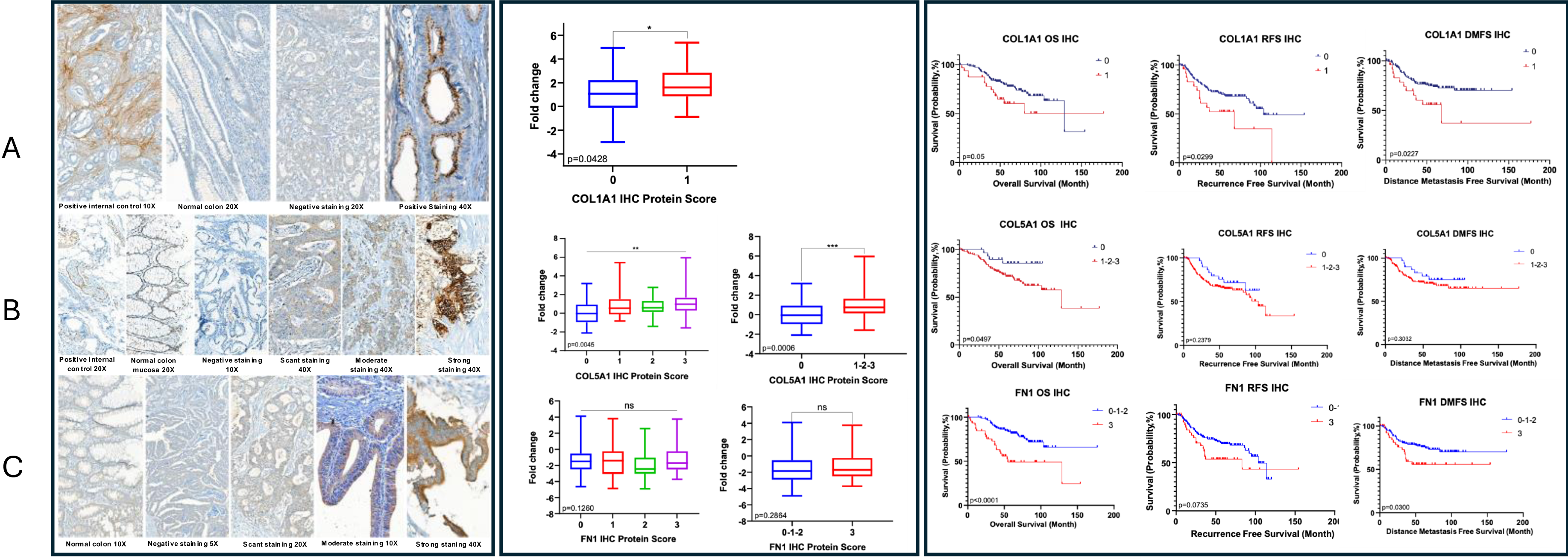
Immunohistochemical (IHC) staining and survival analyses of COL1A1, COL5A1, and FN1 in the retrospective (discovery) cohort. (A) COL1A1 IHC staining showed a predominantly cytoplasmic pattern with varying intensities. (B) COL5A1 IHC staining was observed in both cytoplasmic and membranous patterns. (C) FN1 IHC staining exhibited a primarily nuclear localization. Box plots show the distribution of mRNA expression levels according to IHC scores. Kaplan-Meier survival curves were generated for overall survival (OS), recurrence-free survival (RFS), and distant metastasis-free survival (DMFS). Statistical significance was assessed using the log-rank test. A. COL1A1, B. COL5A1, C. FN1.

COL5A1 IHC staining intensity was also evaluated semiquantitatively by a pathologist and categorised into four scores: 0 (negative), 1 (scant), 2 (moderate), and 3 (strong). Scant and moderate staining were primarily cytoplasmic, while strong staining displayed both cytoplasmic and membranous patterns. Positive internal controls validated the staining procedure. A significant positive correlation was found between COL5A1 protein expression (IHC score) and mRNA expression levels (Figure 4B, left and middle panels). Although patients with positive COL5A1 staining tended to have shorter overall survival, this difference was insignificant. Similarly, no significant association was observed between COL5A1 staining and recurrence-free or metastasis-free survival (Figure 4B, right panel).

FN1 IHC staining intensity was assessed in four categories: 0 (negative), 1 (scant), 2 (moderate), and 3 (strong). Scant and moderate staining was predominantly cytoplasmic, while an apical/luminal pattern characterised strong staining. A significant correlation was observed between FN1 protein expression (IHC score) and mRNA expression levels (Figure 4C, left and middle panels). Survival analysis revealed that patients with strong FN1 staining (score 3) had significantly shorter overall and metastasis-free survival compared to those with lower staining intensities (scores 0-2) (Figure 4C, right panel).

### COL1A1, COL5A1, FN1 and THBS2 Expression Levels in Colorectal Cancer (CRC) Cases’ Serum and their Association with the Disease Prognosis

Another aim of this study was to determine the possible relationship between serum levels of the four genes, whose usability as prognostic biomarkers was investigated, and the disease in CRC patients. For this purpose, the levels of COL1A1, COL5A1, FN1 and THBS2 proteins were determined in the sera of the cases in the prospective validation (prospective) cohort by ELISA.

Accordingly, it was shown that the amount of COL1A1 was statistically significantly higher in metastatic cases compared to non-metastatic cases, and in late-stage cases (stage 3 and 4) compared to early-stage cases (stage 1 and 2) (Figure 5A 1st and 2nd columns). In addition, it was determined that the COL1A1 level was significantly higher from stage 1 to stage 2 and from stage 3 to stage 4 (Figure 5A 3rd column). It was found that the COL1A1 level did not differ between grades (Figure 5A 4th column).

**Figure 5.**
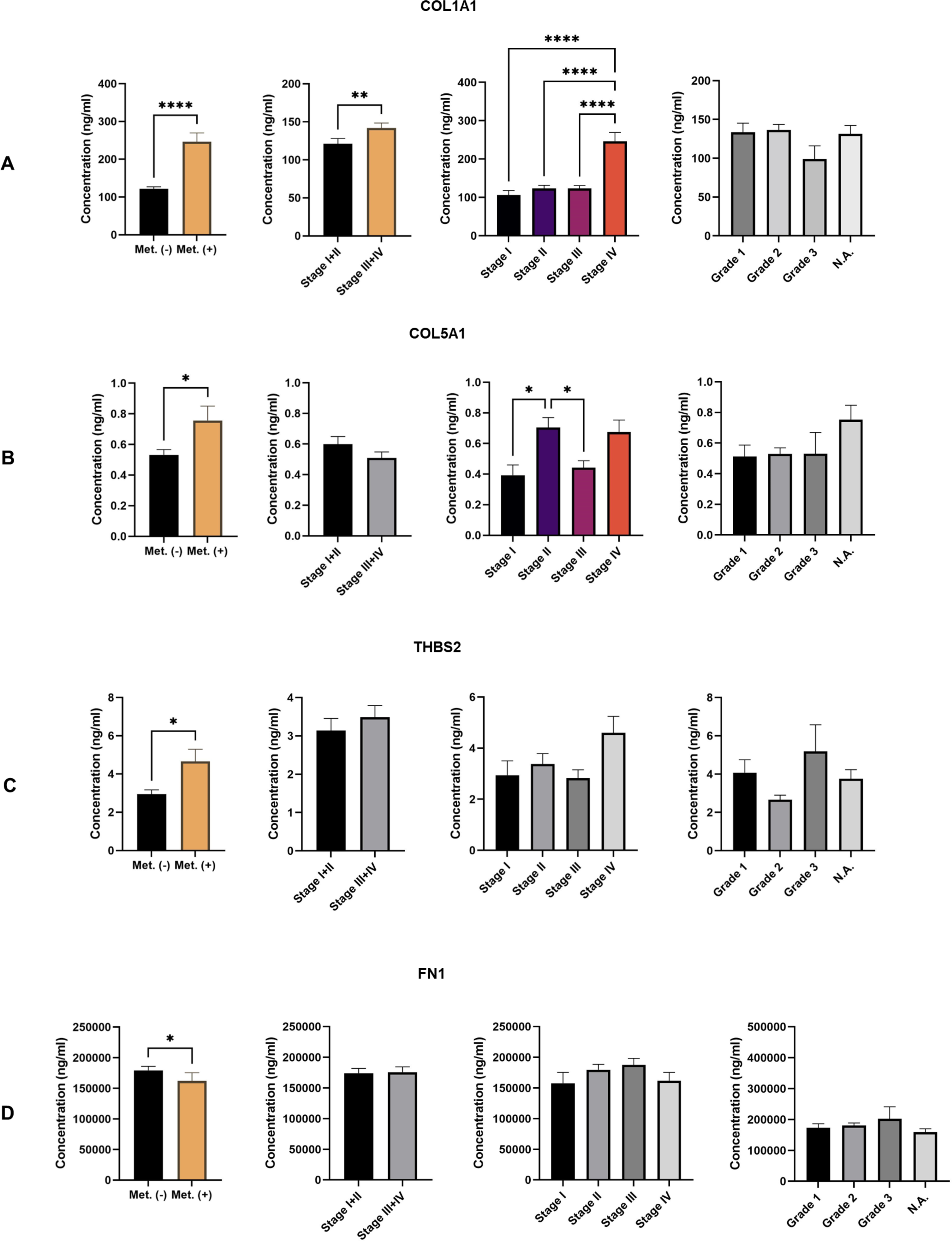
Serum levels of COL1A1, COL5A1, THBS2, and FN1 in the prospective (validation) cohort according to metastasis status, stage, and grade. Bar graphs show each gene’s mean serum protein levels (ng/ml). (A) COL1A1 levels in metastatic vs. non-metastatic cases, and across different stages. (B) COL5A1 levels in metastatic vs. non-metastatic cases, and across different stages. (C) THBS2 levels in metastatic vs. non-metastatic cases. (D) FN1 levels in metastatic vs. non-metastatic cases. Statistical significance was assessed using the t-test or ANOVA. *p < 0.05, **p < 0.01, *p < 0.001.

While it was shown that the amount of COL5A1 was significantly different in metastatic cases compared to non-metastatic cases, a significantly higher amount of COL5A1 was detected in stage 2 cases compared to stage 1 cases (Figure 5B 1st column). An unexpected finding was that stage 2 cases showed higher COL5A1 levels than stage 3 (Figure 5B 3rd column).

Unlike the mRNA data of the discovery (retrospective) and validation (prospective) cohorts, the amount of FN1 was significantly higher in non-metastatic cases compared to metastatic cases (Figure 5C 1st column).

It was shown that the amount of THBS2 was significantly higher in metastatic cases compared to non-metastatic cases (Figure 5D 1. Column).

## DISCUSSION

The need for novel prognostic biomarkers in colorectal cancer (CRC) is underscored by the disease’s high prevalence and mortality rates, coupled with the limitations of existing diagnostic and prognostic tools. Recent studies have identified various potential biomarkers across different biological matrices, including blood, stool, and tissue samples. For instance, circulating exosomal microRNAs, have shown promise as non-invasive diagnostic and prognostic markers for CRC [21].

In this study, we showed that COL1A1, COL5A1, FN1, and THBS2 genes’ mRNA level were higher in tumour tissue samples compared to normal tissue and also higher in metastatic cases compared to non-metastatic cases in our discovery (retrospective) and validation (prospective) CRC cohorts and in recurrent cases compared to non-recurrent cases in retrospective CRC cohort.

When the obtained results are evaluated in detail for each gene, it was shown that COL1A1, one of the genes that play a critical role in the epithelial-mesenchymal transition, which is critical for the cancer metastasis process [22].This study by Zhang et al. showed that COL1A1 gene expression was high at the mRNA level in 24 paired CRC tumour-normal tissue samples. In addition, on a commercial tissue microarray consisting of 75 CRC tumour-healthy tissue, COL1A1 was found to be high in protein expression. It was associated with serosal invasion and lymphatic metastasis. Another study on a microarray dataset including 98 tumours and matched normal tissues determined that high COL1A1 expression levels could predict relapse in high-risk stage II and stage III CRC patients [11]. COL1A1 was one of the 93 differentially expressed proteins identified by Zou et al. in their proteomic analysis of serum samples from 91 CRC cases at different stages. This study, which found that the expression level of COL1A1 was high in tumour samples compared to matched normal tissues at the mRNA level, showed that COL1A1 was high in stage 1 and 2 patients compared to stage 3 and 4 patients and that this situation was correlated with western blot results. In the second sample group included in the study, which consisted of serum samples of 124 independent cases, the concentrations of COL1 and its degradation products PICP and CTx were examined by ELISA. As a result, no difference was found between healthy and cancer samples at the COL1 level. However, the concentration of CTx and PICP was found to increase as the stage increased [23]. In a study supporting the results of this study, collagen type I, III and IV degradation fragments (CIM, C3M and C4M) and type III collagen formation (Pro-C3) levels were measured by ELISA in colorectal cancer, colorectal adenoma carriers and healthy controls. As a result, it was shown that CIM, Pro-C3 and C3M levels were higher in CRC patients compared to adenoma carriers and healthy individuals (CIM, Pro-C3) and that all four biomarkers could distinguish metastatic cases from non-metastatic ones [24].

The present study showed that COL1A1 mRNA was expressed at higher levels in 130 CRC tumour tissues vs 70 unmatched normal colon tissues and 160 CRC tumour-matched normal tissues. Cases with high COL1A1 expression levels have shorter RFS and DMFS (Fig 3A). In IHC analysis, it was also found that patients with high COL1A1 expression levels showed shorter survival in RFS and DMFS (Fig 4A). Moreover, in the ELISA analysis of the serum samples of the prospective cohort, in parallel with Kehlet et al.’s C1M results, it was observed that COL1A1 expression was higher in metastatic cases compared to non-metastatic cases and in advanced-stage cases compared to early-stage cases. These findings indicate that COL1A1 can be used as a non-invasive biomarker in the prognostic follow-up of CRC.

Regarding COL5A1, which is the second focus of this study, there are publications in the literature on different cancer types indicating that this gene is expressed at higher levels in both mRNA and protein levels in tumour tissue compared to normal tissue and that this expression level is associated with disease stage, recurrence and metastasis. In this context, it has been determined that COL5A1 has a prognostic biomarker feature in ovarian cancer [13] (Zhang, 2021), glioma [25], breast cancer [26], tongue squamous cell carcinoma [27] and clear cell renal carcinoma [28]. A bioinformatic analysis performed in colorectal cancer showed that high expression of COL1A1, COL5A1 and COL5A2 correlates with the prognosis of the disease [29].

Another pan-cancer study showed that COL5A1 was highly expressed in 13 cancers, including colorectal cancer, and that this expression was correlated with the prognosis of the disease in 11 cancers. This study, which started with a comprehensive bioinformatic analysis, was also validated by IHC and western blot analysis on five tumour samples and three normal tissue samples representing eight cancer types, including colon cancer [14].

A limited number of studies investigating the presence of COL5A1 in serum as a biomarker candidate. One of them is the study by Nissen et al. investigating the propeptide type V collagen (PRO-C5) level in pancreatic ductal carcinoma (PDAC). This study found that the level of PRO-C5 was higher in the discovery (retrospective) and validation (prospective) cohorts compared to healthy controls and that high PRO-C5 levels were associated with shorter OS, especially in stage II patients. It was found that PRO-C5 level was expressed higher in 8 of the 10 solid tumour types, including colorectal tumours, included in this study with 20 representative samples [30].

In this study, COL5A1 mRNA levels were significantly higher in tumour tissues than in normal tissues in both discovery (retrospective) and validation (prospective) cohorts (Fig 1B). In the discovery cohort, COL5A1 mRNA levels were higher in metastatic cases than non-metastatic cases (Fig 2B). In the same cohort, COL5A1 levels were determined to be higher in patients who died than in those who survived and in those with recurrence than in those without. On the other hand, recurrence-free survival (RFS) and distant metastasis-free survival analyses showed that high COL5A1 levels reduced recurrence-free survival and distant metastasis-free survival in patients (Fig 3B). IHC analyses performed in the discovery cohort showed that the mRNA results in the same cohort were correlated with IHC protein scores. However, COL5A1 IHC results were not associated with OS, RFS and DMFS (Fig 4B). In the validation (prospective) cohort, COL5A1 was significantly higher in metastatic tumour (liver) tissues than in normal tissues of metastatic cases. In addition, it was found that COL5A1 levels were higher in metastatic tumours than in primary tumours in the same cohort (Fig 2B). Serum analyses in the validation (prospective) cohort showed higher COL5A1 in the serum samples of metastatic cases than in the serum samples of non-metastatic cases.

On the other hand, while a higher COL5A1 level was detected in stage II cases compared to stage I cases in the same patient group, a lower amount of COL5A1 was detected in stage III cases compared to stage II cases. It was also observed that the amount of COL5A1 was independent of grade (Fig 5B). However, to the best of our knowledge, this is the first study to comprehensively evaluate the prognostic role of COL5A1 in colorectal cancer (CRC) by integrating biological and bioinformatics analyses, assessing its expression in both tumour and metastatic tissues, as well as at the RNA and protein levels in both tissue and blood samples.

The third ECM gene investigated in this study as a promising biomarker was THBS2. The literature contains significant findings indicating that THBS2 may be a prognostic biomarker for CRC. In the studies of Wang et al., high expression of THBS2, which was identified as one of the genes associated with recurrence in a CRC transcriptome dataset, was shown to be associated with poor prognosis and low survival at both mRNA (QRT-PCR) and protein (IHC) levels, first in independent datasets and then in a group of CRC tumour samples [31]. A study in which a transcriptomic and proteomic integrative meta-analysis was performed showed that high expression of BGN and THBS2 genes was associated with survival, and the effects of these two genes on invasion and migration were also shown on colorectal cancer cell lines ([16]He, 2022). In the studies of Qu et al., it was shown by QRT-PCR and IHC that THBS2 is expressed at higher levels in CRC tissue compared to normal colon tissue and that this situation is associated with low survival [32].

The serum level of THBS2 as a potential biomarker has been investigated in non-small-cell lung cancer [33], pancreatic cancer and distal cholangiocarcinoma [34] and gastric cancer [35], and it has been shown that high THBS2 expression can distinguish cancer cases from healthy controls. In the study by Fei et al., the level of THBS2 was examined together with RBP4 in CRC patients. In this analysis, which was performed in the serum of 402 CRC patients and 218 healthy control individuals, it was shown that the expression level of THBS2 was higher in healthy control individuals compared to CRC patients and that the two groups could be distinguished accordingly [36].

This study showed that THBS2 was expressed at higher levels in tumour tissues than in normal colon tissues at the mRNA level in both discovery (retrospective) and validation (prospective) cohorts (Fig 1C). On the other hand, it was found that THBS2 was higher in metastatic cases than in non-metastatic cases at the mRNA level in the discovery (retrospective) and validation (prospective) cohorts (Fig 2C). In the discovery (retrospective) cohort, THBS2 gene expression was higher in deceased cases than in survivors and recurrent cases compared to non-recurrent cases. On the other hand, THBS2 expression level was not associated with RFS and DMFS (Fig 3C).

Finally, regarding FN1, which is involved in ECM remodelling, some studies associate it with poor prognosis and metastasis in non-small-cell lung cancer, head and neck cancer, breast cancer, nasopharyngeal cancer and gastric cancer [15, 37–40]. There are studies with different findings on the prognostic potential of FN1 in colorectal cancer. It has also been shown that silencing of FN1 suppresses the proliferation, migration and invasion of colorectal cancer [41]. In this study by Cai et al., it was shown by IHC and QRT-PCR that FN1 was expressed at higher levels in tumour tissues compared to matched normal tissues, and the relationship between FN1 and prognosis was determined using TCGA data. In the study conducted by Bogdanovic et al. on liver metastasis tissue and matched normal tissue and serum samples of 30 colorectal liver metastasis patients, FN1 was not associated with recurrence, and FN1 expression was found to be higher in liver normal tissue compared to metastatic tumour tissue [42]. IHC and QRT-PCR analysis performed in 107 CRC tumour tissues and 115 normal colon tissues that were not matched showed that FN1 was expressed at higher tumour tissue levels than the normal colon at both mRNA and protein levels. Overall survival analysis in this patient group indicated that patients with high FN1 expression had lower survival [43]. In a study analysing Extra Domain A (EDA), one of the alternative splices of FN1, in the sera of 70 colorectal cancer patients, it was determined that EDA level was higher in CRC patients compared to healthy individuals and in late-stage patients compared to early-stage patients. The same study also showed that high EDA levels shortened the overall survival of CRC patients [44]. In a proteomic analysis and immunoblotting study performed in the plasma of 10 metastatic, 10 non-metastatic CRC patients and 10 healthy control individuals, it was shown that FN1 was lower in CRC patients compared to healthy controls, and this was shown in both the discovery (n=30) and validation (n=26) cohorts of this study [45]. A proteomic study performed on extracellular vesicles isolated from plasma samples of CRC patients showed that high expression of FN1 was correlated with the progression of CRC [46].

In this study, while FN1 was expressed at higher levels in CRC tumours than in normal colon tissues in the discovery (retrospective) cohort, the opposite was found in the validation (prospective) cohort (Fig 1D). Similarly, while higher FN1 levels were observed in metastatic CRC patients compared to non-metastatic ones in the discovery (retrospective) cohort, higher FN1 expression was detected in matched normal liver tissues of metastatic cases compared to metastatic liver tumour tissues, in line with Bogdanovic et al.’s study. In the analysis of the patients in the discovery (retrospective) cohort, while higher FN1 expression was observed in patients with recurrence compared to those who did not have recurrence, FN1 expression was not associated with survival (Fig 3D). IHC analysis with FN1 did not show a significant difference between the increase in FN1 protein expression and mRNA expression. FN1 IHC results were not associated with survival (Fig 4D). Analysis of serum samples from the validation (prospective) cohort showed that FN1 levels were higher in non-metastatic patients compared to metastatic patients. Still, there was no significant association between CRC stage or grade increase and FN1 levels (Fig 5D).

Controversial roles have been attributed to FN1 in cancer progression. Fibronectin has long been recognised as an acute-phase reactant synthesised by hepatocytes in response to various inflammatory processes. The soluble plasma fibronectin isoform lacks the EDA and EDB modules and mainly originates from hepatocytes. In contrast, extracellular matrix fibronectin includes an EDA segment which regulates cell cycle progression. This domain acts as a mitogenic signal and promotes epithelial-mesenchymal transition [47–51]. Therefore, discriminating between the soluble and ECM isoforms of FN1 may offer a more accurate prognostic assessment.

## CONCLUSION

In our previous study (Belder et al.), we validated four ECM genes with prognostic biomarker potential, which we obtained as a result of the integration of the transcriptome data we obtained from 49 paired CRC tumour-normal tissues and the GEO datasets containing 251 CRC tumor-50 normal colon tissue transcriptome, in an independent cohort including 64 independent paired CRC tumour and normal colon tissues. Based on the expression levels of these four genes, we developed a prognostic risk score model that successfully stratified CRC patients into high-risk and low-risk groups, with the high-risk group showing significantly poorer overall survival (OS) and more unfavourable clinicopathological features. In the present study, we analysed these four genes at the mRNA level in a retrospective discovery (retrospective) cohort consisting of 130 CRC (including 64 CRC cases included in the previous study) and 70 normal colon tissues and a validation (prospective) cohort consisting of 160 paired CRC tumours and normal colon tissues, at the protein level by IHC in the discovery (retrospective) cohort tumour tissue, and by ELISA in the validation (prospective) cohort, which included the serum of 180 CRC patients. Our results revealed that COL1A1, COL5A1 and THBS2 were higher in both primary and metastatic tumour tissues compared to normal tissues at the mRNA level and higher in metastatic cases compared to non-metastatic cases at the serum protein level. The high level of FN1 mRNA expression could not be confirmed in our validation (prospective) cohort. IHC analyses showed that high expression of COL1A1 and FN1 reduced survival. In this study, it was shown for the first time in the literature that COL1A1 and COL5A1 proteins were expressed at higher levels in the serum samples of a large CRC cohort in metastatic cases compared to non-metastatic cases and that this was correlated with the tissue mRNA profile of the same patient group. In conclusion, our study provides novel insights into the prognostic significance of COL1A1, COL5A1, and THBS2 as biomarkers for CRC, particularly in metastatic cases. By integrating both mRNA and protein expression levels, including serum protein analysis, we have demonstrated the potential of these genes to serve as diagnostic and prognostic markers for CRC, with implications for personalised treatment strategies and improved patient management.

## MATERIALS & METHODS

### Patient Cohorts and Sample Preparations

Our previous colorectal cancer (CRC) transcriptome profiling study identified four candidate genes and demonstrated their association with CRC development, progression, and poor prognosis. Building upon these findings, the current study employed two independent cohorts— the discovery (retrospective) cohort and the validation (prospective) cohort—to further evaluate the biomarker potential of these four genes in CRC. These cohorts were explicitly designed to assess the biomarkers in both primary and metastatic CRC cases and their potential as blood-based diagnostic markers.

### Discovery (Retrospective) Cohort

The discovery (retrospective) cohort included 200 colorectal cancer tumour tissues, of which 49 were metastatic cases and 70 matched adjacent normal tissues collected from the same patients. Matched normal tissues were carefully selected from the furthest region of the surgical margin to minimise the risk of tumour involvement. This cohort comprised archived samples from patients diagnosed with colorectal adenocarcinoma at the Ankara University Faculty of Medicine between 2006 and 2015. Formalin-fixed, paraffin-embedded (FFPE) blocks used for this cohort were 2–8 years old. Hematoxylin and eosin (H&E)-A pathologist reviewed stained slides from each case to identify tumor-enriched areas. Four 8-μm thick sections were cut from each FFPE block, with two sections placed on each standard glass slide. One tissue section was stained with H&E to guide the identification and marking of tumour regions. Macrodissection of the marked tumour regions was performed on serial sections for RNA extraction. Immunohistochemistry (IHC) analyses were conducted on tumour tissues from all 200 patients to validate the protein-level expression of four target genes.

### Validation (Prospective) Cohort

The validation (prospective) cohort included 160 paired primary colorectal cancer tumor and normal tissues, 25 of which were obtained from metastatic cases. These fresh-frozen samples were collected during surgery and immediately preserved in cryotubes containing QIAGEN RNAprotect Tissue Reagent (Cat. No. 76106) following the manufacturer’s protocol.

Additionally, peripheral blood samples were collected from 180 patients, including those in the validation (prospective) cohort, to evaluate the diagnostic potential of the four target genes at the protein level using enzyme-linked immunosorbent assay (ELISA).

### Ethical Approval

Patient characteristics for both the discovery (retrospective) and validation (prospective) cohorts are summarised in Table 1. Ethical approval for the discovery (retrospective) cohort was obtained from the Ankara University School of Medicine Clinical Research Ethics Committee (Ref: 153-4854), and approval for the validation (prospective) cohort was granted by the Ankara Eğitim Araştırma Hospital Ethics Committee (Ref: 15-200-19). Written informed consent was obtained from all participants. The study adhered to the principles outlined in the Declaration of Helsinki.

### RNA extraction and quantitative RT-PCR

For FFPE samples from the discovery (retrospective) cohort, RNA was isolated from the marked, macrodissected tumour regions using the RNeasy FFPE kit (Qiagen, Hilden, Germany), with a modified deparaffinisation procedure as described in our previous study (Belder et al., 2016). Fresh frozen tissue samples were thawed and placed on dry ice for the discovery (retrospective) cohort. Tissue disruption and homogenisation were performed using the QIAGEN TissueLyser LT device. Following homogenisation, RNA was isolated using the RNeasy kit (Qiagen, Hilden, Germany) according to the manufacturer’s protocol. The quantity and purity of all isolated RNA samples were assessed using a NanoDrop ND-1000 Spectrophotometer.

cDNA synthesis was performed from 0.5 μg of total RNA using the Transcriptor First Strand cDNA Synthesis Kit (Roche Diagnostics, Basel, Switzerland), following the manufacturer’s instructions. Real-time PCR was then conducted using the SYBR Green I Master kit (Roche Diagnostics, Basel, Switzerland) on a LightCycler480 instrument for quantitative expression analysis, with triplicate measurements of each sample. HPRT and B2M were employed as housekeeping genes. The data were analyzed using the ΔΔCT method (Livak and Schmittgen, 2001). The sequences of the qRT-PCR primers used in this study are provided in Supplementary Supplementary Table 1.

### Assessment of the Association Between Expression Patterns of THBS2, FN1, COL1A1, and COL5A1 and Recurrence and Metastasis in Colorectal Cancer Using Our Cohorts and Public Datasets

The association between high gene expression of THBS2, FN1, COL1A1, and COL5A1 and advanced tumour progression (including recurrence and metastasis) was assessed using our discovery cohorts. The correlation between the expressions of these four genes and clinical characteristics, including recurrence, metastasis, and survival outcomes, was analysed. Box plots were used to display the differences in the expression levels of THBS2, FN1, COL1A1, and COL5A1 based on clinical factors, including recurrence status, presence of metastasis, and mortality (alive vs. death). These plots illustrate how the expression of each gene varies with these clinical outcomes. A p-value of <0.05 was considered statistically significant. Kaplan-Meier analysis for relapse-free and distant metastasis-free survival was also conducted using the discovery cohort qPCR results to analyse the correlation between the expression of these four genes and CRC prognosis. The samples were divided into high- and low-expression groups, and the results were assessed using Kaplan-Meier survival plots.

### Immunohistochemistry analysis

Immunohistochemistry analysis was performed on FFPE tumour tissue sections from 200 patients (discovery cohort) using a Ventana BenchMark ULTRA platform (Ventana Medical Systems, Tucson, Arizona). The following primary antibodies were used: FN1, (1:100) (ab6328, Abcam, UK); COL1A1, (1:200) (ab138492, Abcam, UK); COL5A1, (1:1000) (HPA030769, Sigma, USA); and THBS2, (1:100) (SAB4301622, Sigma, USA). Four μm-thick sections from each macroarray block were mounted on slides, labelled with unique barcodes containing protocol information, and incubated at 70°C for 60–120 minutes for paraffin melting and tissue adhesion. After manual application of primary antibodies and incubation, slides were processed using the UltraView Universal DAB Detection Kit, which includes a secondary antibody conjugated to horseradish peroxidase (HRP), producing brown precipitates upon reaction with hydrogen peroxide and DAB chromogen. An expert pathologist evaluated the slides, scoring staining intensity (0: none, 1: weak, 2: moderate, 3: strong) based on the percentage, intensity, and localisation of staining (membranous, cytoplasmic, etc.).

### ELISA

FN1, COL1A1, THBS2, and COL5A1 protein levels in serum samples (n=180) were determined using ELISA. Each analysis was performed in duplicate and according to the manufacturer’s protocol. The Abcam Human Fibronectin ELISA Kit (Cat. No: ab219046) was used for FN1, and the Abcam Human Pro-Collagen I alpha 1 ELISA Kit (Cat. No: ab210966) for COL1A1 (Detection range: 0.125-8 ng/ml and 3.906-2.5 ng/ml; Sensitivity: 2.006 ng/ml and 0.053 ng/ml, respectively). The Abnova THBS2 Human ELISA Kit (Cat. No: KA0667) was used for THBS2 (Detection range: 0.625-40 ng/ml; Sensitivity: 0.01 ng/ml). The CUSABIO Human Collagen, type V, alpha 1 (COL5A1) ELISA Kit (Cat. No: CSB-E13447h) was used for COL5A1 (Detection range: 0.312-20 ng/ml; Sensitivity: 0.078 ng/ml). Plate readings were performed at 450 nm using the TECAN INFINITE M PLEX device. Protein levels in serum samples were calculated using linear regression based on standard O.D. values for FN1 and COL1A1 and using a 4PL regression model for THBS2 and COL5A1.

### Statistical analysis

Statistical analysis was performed using GraphPad Prism (version 8.3.0, GraphPad Software Inc., San Diego, CA, USA) and IBM SPSS® Statistics (version 20.0; IBM Company, New York, NY, USA). Statistical significance was defined as p < 0.05 for all analyses. Unpaired Student’s t-tests were used for normally distributed data, and non-parametric Mann-Whitney U-tests were used for non-normally distributed data to compare central tendencies. Significance was set at p < 0.05. To assess relapse-free survival, metastasis-free survival, and overall survival, Kaplan-Meier survival analysis with a log-rank test was performed. The median expression of the genes was used as the cutoff value for stratifying the patients. Furthermore, univariate and multivariate Cox regression analyses were performed to evaluate the prognostic relevance of the four gene expressions in CRC patients. Hazard ratios (HRs) and 95% confidence intervals (CIs) were calculated.

## Supporting information

Sequences of the primers used of QRT-PCR

## Acknowledgement

This study is supported by Ankara University BAP project no. 20B0415001, TÜBİTAK TEYDEB project no. 3210467.

