## Supplementary material for "Evaluation of the Prognostic Value of Four ECM-Associated Genes in Recurrence and Metastasis of Colorectal Cancer": Sequences of the primers used of QRT-PCR

| **Genes** | **Forward Primer (5’—3’)** | **Reverse Primer (5’—3’)** | **Amplicon length** | **Annealing Temperature** |
| --- | --- | --- | --- | --- |
| FN1 | GCTCTATTCCACCTTACAACAC | ACACAACGATGCTTCCTGAG | 155 | 55 |
| COL1A1 | AAGAGGAAGGCCAAGTCGAG | AGATCACGTCATCGCACAAC | 156 | 56 |
| COL5A1 | GATGTCGCTTACAGAGTCACC | TTGGCTTTCACAGTTGTTAGGA | 112 | 55 |
| THBS2 | AGCTCAGCGAGAACCTCAAG | TTCATTTTCCGCAAAGAACC | 128 | 60 |
| B2M | TGCTGTCTCCATGTTTGATGTATCT | TCTCTGCTCCCCACCTCTAAGT | 86 | 57 |
| HPRT | TGACACTGGCAAAACAATGCA | GGTCCTTTTCACCAGCAAGCT | 94 | 59 |
